# A Cretaceous bug with exaggerated antennae might be a double-edged sword in evolution

**DOI:** 10.1101/2020.02.11.942920

**Authors:** Bao-Jie Du, Rui Chen, Wen-Tao Tao, Hong-Liang Shi, Wen-Jun Bu, Ye Liu, Shuai Ma, Meng-Ya Ni, Fan-Li Kong, Jin-Hua Xiao, Da-Wei Huang

**Affiliations:** Institute of Entomology, College of Life Sciences, Nankai University, Tianjin 300071, China; Key Laboratory of Zoological Systematics and Evolution, Institute of Zoology, Chinese Academy of Sciences, Beijing 100101, China; Beijing Forestry University, Beijing 100083, China; Paleo-diary Museum of Natural History, Beijing 100097, China; Fujian Paleo-diary Bioresearch Centre, Fuzhou 350001, China; Century Amber Museum, Shenzhen 518101, China

**Author notes:** These authors contributed equally (Bao-Jie Du, Rui Chen). Correspondence and requests for materials should be addressed to D.W.H. or J.H.X.

**Keywords:** Cretaceous, Coreidae, sexual selection, defense, costs

## Abstract

In the competition for the opposite sex, sexual selection can favor production of exaggerated features, but the high cost of such features in terms of energy consumption and enemy avoidance makes them go to extinction under the influence of natural selection. However, to our knowledge, fossil on exaggerated traits that are conducive to attracting opposite sex are very rare. Here, we report the exaggerated leaf-like expansion antennae of *Magnusantenna wuae* Du & Chen gen. et sp. nov. (Hemiptera: Coreidae) with more abundant sensory hairs from a new nymph coreid preserved in a Cretaceous Myanmar amber. The antennae are the largest among species of coreid and one of the largest known insects. Such bizarre antennae indicate that sensitive and delicate sensory system and magnificent appearance in Hemiptera have been already well established in mid-Cretaceous. Our findings provide evidence for Darwin’s view that sensory organs play an important role in sexual selection. This nymph with the leaf-like antennae may also represents a new camouflage pattern for defense. However, the oversized antennae are costly to develop and maintain, increasing the risks from predators. Such unparalleled expanded antennae might be the key factor for the evolutionary fate of this Myanmar amber coreid species.

**Significance:** Darwin proposed the importance of sensory organs in sexual selection, but it was greatly ignored compared with weapons and other common ornaments. Here, we report a new type of insect antennae, the multiple segments leaf-like expansion antennae from a new nymph coreid preserved in a Cretaceous Myanmar amber. Our finding provides evidence for the prominent role of sensory organs in sexual selection and thus supports Darwin’s viewpoint. This discovery demonstrates that high-efficiency antennae were present in Coreidae 99 million years ago. In addition, the exaggerated antennae might represent a new evolutionary innovation for defensive behavior. This is a case in which the high benefits and high costs brought by the exaggerated antennae jointly determine the direction of species evolution.

## Introduction

Luxurious and decorative structures are usually regarded as weapons or ornaments, and are vital features in sexual selection, including feathers, antlers, pebbles, horns, jaws, etc. (1–5). Female mate choice and male-male competition further complicate and beautify male specialized characters (6). Just as Darwin once proposed, the success rate of animal mating depends not only on the above two widely accepted mechanisms of sexual selection, but also on the efficient locating to the mate, that is, the sense organ plays an important role in sexual selection (7). Chemical and acoustic signals seem to be most commonly used to locate mates, especially in long distances (8). However, Darwin’s idea has been more or less ignored (6, 9). At present, most studies attribute the mate’s positioning to communication behavior rather than sexual selection behavior (10, 11). There are only a few cases where sensory organs are associated with sexual selection: the size of thoracic spiracles in katydids (12), the length of antennae in moths (6, 13), the number of sensory structures of antennae in false garden mantids (14). The researches on sensory organs mainly focus on the morphology of antennae, expansive and delicate antennae carry more olfactory receptors, which facilitates the location of mate (14, 15). Once an individual enters the process of short distance courtship, visual signals play a significant role, surprisingly, the importance of antennae as an important display function in sexual selection has been greatly overlooked. Fossil materials have provided evidence for the origin and evolution of important characters of many species, as well as related behavioral information (16–19), but the materials related to the functional morphology of specialized antennae are rare. Some insects with ramified antennae exhibited chemical communication in the early Cretaceous, but not show the sexual display behavior (15, 20–22). So far, the only fossil species of Hemiptera with leaf-like dilated antennae are *Reticulatitergum hui* Du, Hu and Yao, 2018 (Yuripopovinidae) from the Cretaceous and *Gyaclavator kohlsi* Wappler et al., 2015 (Tingidae) from the Eocene, and the expansion of their antennae occurs in the fourth antennal segment (23, 24). A few of extant Coreids have the similar antennae (25). However, there is no evidence that the antennae of Cretaceous insects play an important role in both mate positioning and sexual display.

Coreidae is a moderately large family in Hemiptera, with nearly 500 genera and 2200 species (26). Coreids are hemimetabolous herbivore insect, commonly known for their leaf-like expansive antennae and legs (27). Expansion of various body parts plays a significant role in sexual selection and defensive behavior (24, 27–29). So far, only five species of the family Coreidae have been described from the Mesozoic rock impressions, all of which do not have expanded antennae (30). Therefore, the origin, evolutionary diversity and corresponding functions of these early exquisite expansions in Coreidae remain poorly understood.

Here, we report a coreid nymph with exaggerated, expanded antennae from mid-Cretaceous Burmese amber. *Magnusantenna wuae* Du & Chen gen. et sp. nov. represents the first record of the family Coreidae preserved in amber. The leaf-like antennal expansion of the coreids is demonstrated to have existed approximately 99 million years ago. This discovery improves our understanding of coreid biodiversity during the Cretaceous and provides evidence for the prominent role of sensory organs in sexual selection. We then discuss the role of the specialized antennae in defensive behavior, as well as the negative effects on survival.

## Systematic Paleontology

Order Hemiptera Linnaeus, 1758; Family Coreidae Leach, 1815; Subfamily Coreinae Leach, 1815; Genus *Magnusantenna* Du & Chen gen. nov.

### Type species. *Magnusantenna wuae* Du & Chen gen. et sp. nov

(Figs. 1, S1–3)

**Fig. 1.**
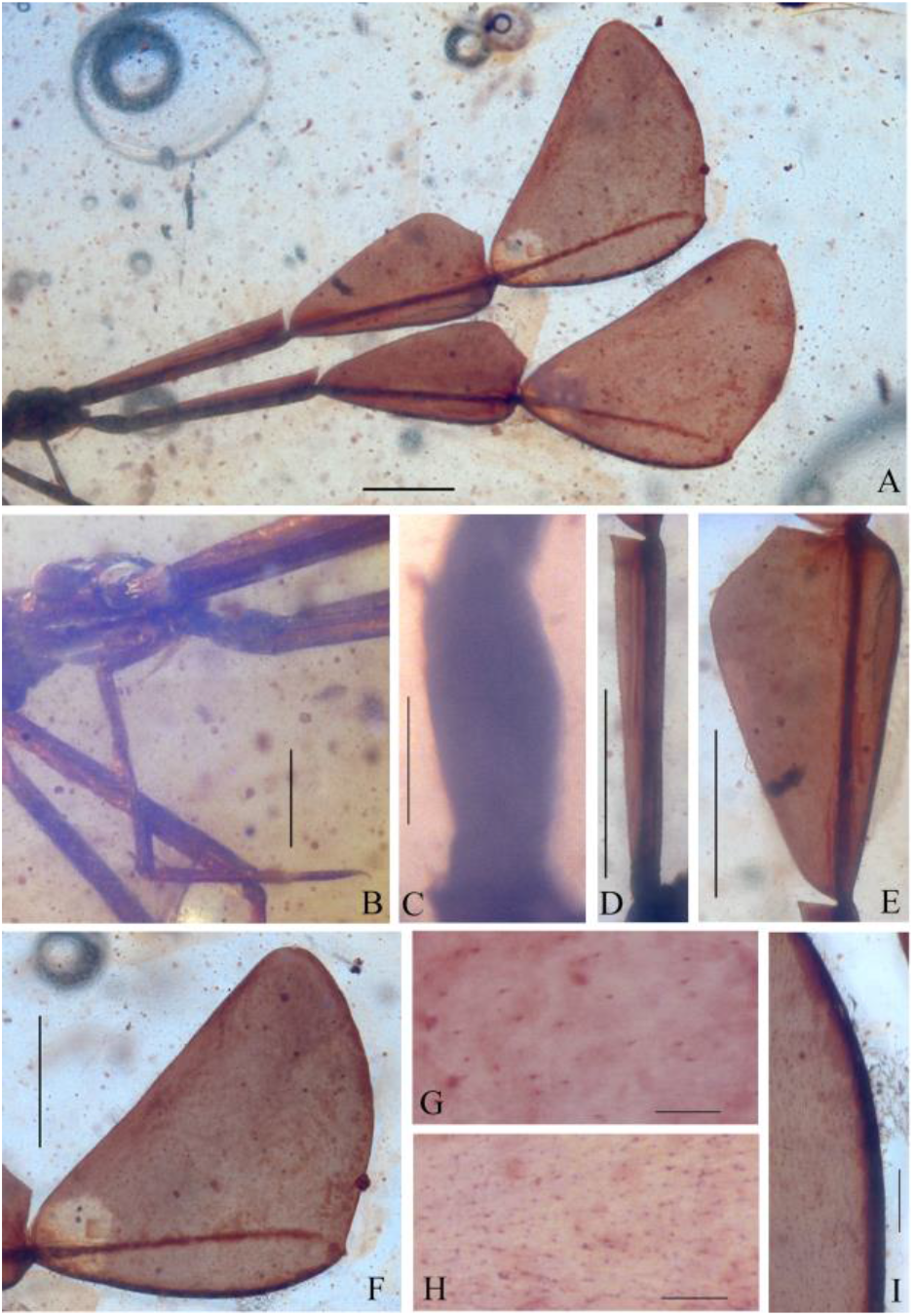
Head of *M. wuae* gen. et sp. nov. (STJS0003). A, C–I. Lateral views of the antenna: (A) Overall view. Scale bar, 1 mm. (C) First segment. Scale bar, 200 μm. (D) Second segment. Scale bar, 1 mm. (E) Third segment. Scale bar, 1 mm. (F) Fourth segment. Scale bar, 1 mm. (G) Setae on the distal expansion of the fourth antennal axis. Scale bar, 100 μm. (H) Setae on the proximal expansion of the fourth antennal axis. Scale bar, 100 μm. (I) Strong keratin thickening of the proximal margin of the fourth antennal axis. Scale bar, 200 μm. (B) Ven-lateral view of head. Scale bar, 500 μm.

#### Etymology

The generic name is derived from Latin prefix *margus*, meaning large, and *antenna*, meaning antenna; referring to the enlarged antennae. The specific epithet, *wuae*, is in honor of Ms. Wu Lijing, who discovered the specimen.

#### Holotype. STJS0003

Nymph, probably approach fourth instar. Gender undeterminable. Deposited in the Century Amber Museum (Room 301A No.1, Songrui Road, Songgang Street, Bao’an District, Shenzhen, China). Only known specimen.

#### Horizon and Locality

Hukawng Village, Kachin State, northern Myanmar; Upper Cretaceous (earliest Cenomanian), 98.79 ± 0.62 Ma (31). Only known from the type locality.

#### Diagnosis

Body slender. Antenna extremely large, subequal to the body length, with four segments. First segment inflated; second, third and fourth segments expanded and remarkably toward the apex. Head square, compound eyes large, spherical, located at the center of each side of the head and prominently protruding. Pronotum and mesonotum trapezoidal. Legs slender. Detailed description provided in SI Text.

## Discussion

### Antennal expansion maintained in adult

Heteropteran nymphs generally have five instars, which typically resemble adults in their morphological appearance and living environment, except that they are generally much smaller and softer than adults, they have paired scent glands located on the dorsal abdomen, and the number of tarsal segments is one less than that of adults (27). Wing buds appear in the third instar, and external genitals and ocelli can be observed in the fifth instar (29). We regard this new specimen as a nearly fourth instar nymph because of the following characters (Fig. 1; see SI for detailed describe; Figs. S1–3): posterior margins of the hind buds not reaching the anterior margin of the first abdominal tergite; ocelli absent; scent glands located on the dorsal surface of the abdomen; tarsi two-segmented; and genitalia not developed (29).

The family Coreidae have the expanded antennae, which are relatively common in living insects and likely to still exist during adulthood (Fig. S4). None of the fossil species of Coreidae has expanded antennae (See SI for detailed fossil records). Both nymphs and adults of modern *Chariesterus, Dalader*, and *Thasus* in Coreinae have the similar expansion on the third antennal segment (25, 32), but are significantly different from the new species with more exaggerated expansion (Fig. 1, Detailed determined of taxonomic status in SI, Figs. S1–3). The first antennal segment of the nymph described here is robust (Fig. 1C, Figs. S1, S3); the proximal expansion of the antennal axis extends from the second to fourth segment with dense setae on the surface, and the proximal margin exhibits strong keratin thickening (Fig. 1A, D–I, Figs. S1, S3); the fourth antennal segment is more prominent than the third segment (Fig. 1A, E–F, Figs. S1, S3). Combining the developmental stage of the nymph and the specialized form of the antennae, we speculate that expanded antennae should continue throughout the adult stage. In addition, we cannot rule out the possibility that adults with larger antennae were not preserved due to the lack of amber’s ability to preserve larger inclusions. The multiple segments expansion antennae represent a new type of insect antennae. This discovery demonstrates that the antennal expansion of Coreidae originated at least 99 million years ago.

### Antennal expansion and sexual selection

Some insect antennae reach remarkable sizes. The antennae in katydids, crickets and some longhorn beetles are often longer than the body, and the antennae in some chafers, moths and mosquitoes are relatively wide in relation to the body, but these antennae cannot be longer and wider than the body at the same time. The antennal length and width of *M. wuae* gen. et sp. nov. preserved in the amber described herein are greater than the body length and width (the width is more than 3 times the body width), which is unique in Heteroptera and rare in insect. The presence of nymph suggests that if we find an adult in the future, the antennae may be larger and expansion of the third and fourth segments of the antennae may be more significantly. The similar large, exquisite lamellate antennae and pectinate antennae significantly increase the surface area of the antennae, and enhance the interaction between odors and receptors (33). When the recognition of certain odors becomes an important factor in reproductive fitness, the evolution of olfactory receptors tends to maximize the surface area of antennae (34). The antennae sensilla capable of obtaining information are mainly located on the distal end and lateral expansion of the flagellum (35, 36). The exaggerated antennae may have borne a large number of olfactory receptors, enabling the coreids to locate mates even when they release relatively low concentrations of pheromones and providing a strong basis for sexual display behavior, improving intraspecific and interspecific competitiveness (15, 22, 35). The leaf-like expansion of the antennae in modern male coreids is usually used to courtship, and the mating success rate is closely related to male body size (24, 27, 37). The special antennae provide a significant visual signal for females to find high-quality males. Normally, only a strong male can support such a large antenna. Therefore, we suspect that the extremely expanded antennae of *M. wuae* gen. et sp. nov. may be used for sexual selection in adult, which is reflected in the locating and attracting to the mates. When the population density is low, the antennae may invest more in positioning females, and when the population density is high, the antennae may invest more in attracting females (9).

In summary, our finding indicates that the coreid has high-efficiency mates positioning and sexual display capabilities during the Cretaceous period. The occurrence of the extremely expanded antennae is a powerful and effective survival strategy. It is the result of natural selection and sexual selection (Fig. 2), and may be strongly influenced by sexual selection (7). This is a case of the emergence of sexual selection characters caused by positive feedback (Fig. 2). Our finding provides evidence for the idea that sexual selection may also act on “organs of sense” suggested by Darwin. Other known Hemipteran fossils with expanded antennae are species from Yuripopovinidae and Tingidae (23, 24). These similar expansions of antennae found in Hemiptera suggest possible behavioral convergence.

**Fig. 2.**
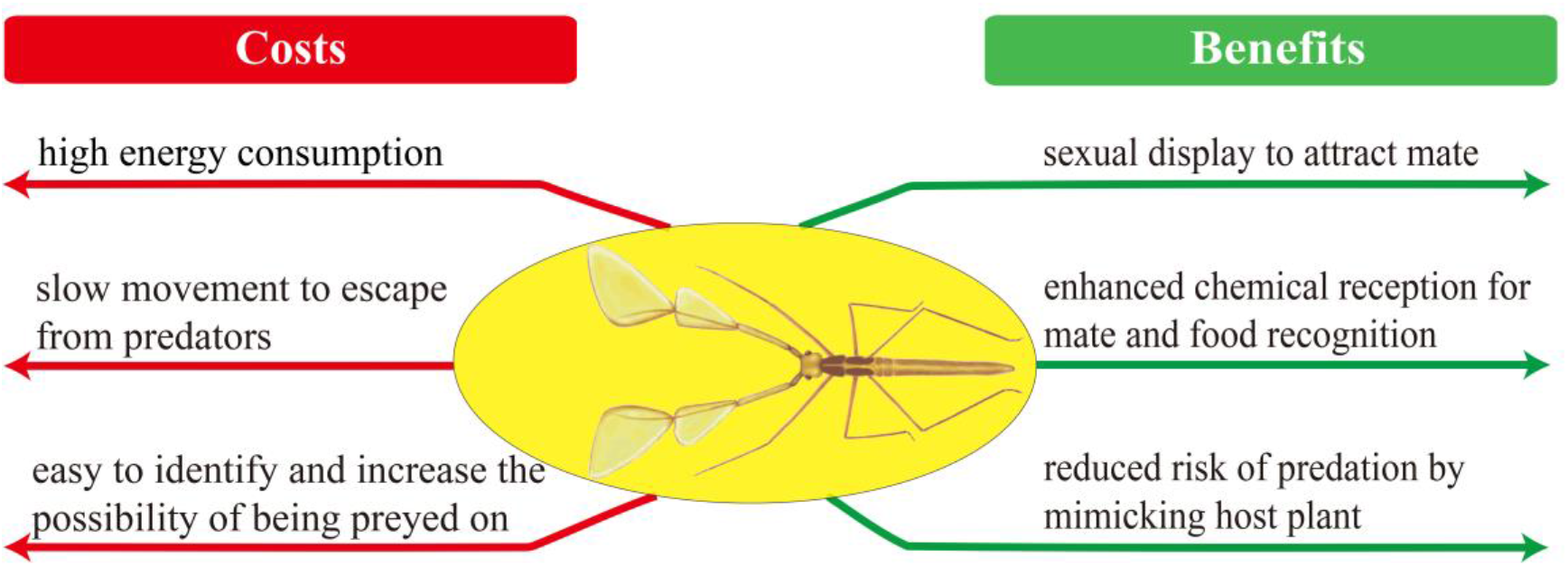
Possible costs and benefits caused by the exaggerated antennae during the evolution of *M. wuae* gen. et sp. nov. (STJS0003).

### Antennal expansion and defensive behavior

Camouflage is a common behavior in insects and can effectively reduce the probability of an individual being preyed by predators. Leaf-like expansion is usually related to the leaves of mimic plants, however, the beneficial behavior in fossils is extremely rare (38–40). Insects have evolved various ways of avoiding predators, but most have focused on the adaptive evolution of their wings (38).

For the nymph described here, it’s probably simulating a branch with leaves in stationary, which is more complex than simply imitating a leaf or a branch. This is a unique and effective defense pattern evolved in response to the numerous predators cotemporaneous, such as reptiles, mammals, birds and other vertebrates, as well as spiders and other insectivorous invertebrates. It provides evidence for the evolution of insect behavior diversity under natural selection (Fig. 2).

Unlike the previous leaf-like mimicry that present on the imaginal wing and the larval thorax and abdomen, the nymph’s leaf-like expansion on the antennae is a new evolutionary innovation. This discovery may represent a new camouflage pattern. It is the earliest record of camouflage behavior in Hemiptera.

### Antennal expansion and possible costs

The balance between the considerable benefits and the enormous costs brought by special features is one of the most exciting and under-researched topics in sexual selection studies (41). Although in some cases, the presence of sexual selection makes adaptation more effective, but the experiments on wild populations have found that sexual selection could restrict the ability of individuals and populations to adapt to changed environments and accordingly increase the risk of species extinction (42–45).

Since the Cretaceous, the rise of herbivorous and carnivorous insects, birds and other animals improved the diversity and number of competitors and predators of the nymph (46, 47). Therefore, the survival pressure of the nymph increased dramatically. As far as the exaggerated antenna is concerned, it requires a lot of energy to produce and maintain, which may need to put more efforts when the competitive pressure increases. Moreover, the spectacular antennae are more likely to have exposed the individual, increasing the chance of being discovered and preyed upon. In addition, the large appendages probably made it move slowly, which would have been a disadvantage when fleeing a predator. The combination of these factors may increase the cost caused by the specialized character in an all-round way (Fig. 2). Similar examples are the exaggerated pod-like tibiae of the dancing dragonfly and the extremely elongated abdomens and sexual organs of the mecopterans (48, 49).

## Material and Methods

The coreid nymph described herein is preserved in a piece of golden-brown Myanmar amber from an amber deposit in the Hukawng Valley of Myanmar. The age has been estimated to be ca. 99 Ma (98.8 ± 0.6 Ma; earliest Cenomanian, Upper Cretaceous) based on U-Pb dating of zircons from the volcaniclastic matrix of the amber-bearing deposit (31). The mining locality is at Noije Bum, near Tanai Village (26°21’33.41”N, 96°43’11.88”’E) (50, 51). Details of the geology and stratigraphy of the deposit have been described in previous publications (31, 50). The piece of amber was cut, ground and polished to a length × width × height of approximately 26.75 × 20.14 × 12.43 mm. The specimen was examined with a LEICA M125 C dissecting microscope. Photographs were obtained with a LEICA MC 190 HD fitted to a LEICA M125 C stereomicroscope and a Nikon Digital Sight DS-Ri1 fitted to a Nikon AZ100M stereomicroscope. Images were stacked with Helicon Focus 6. Photographic figures were constructed in Adobe Photoshop CC.

## Acknowledgements

We are grateful to Century Amber Museum for depositing the specimen. We sincerely thank Max Barclay at the Natural History Museum in London for his valuable comments on our article. We would like to express our gratitude to Ms. Melanie Schuchart, Mr. Steve Kerr, Mr. Branco, Kwan and Dr. Margarethe Brummermann for providing us with photos. This project is supported by the National Natural Science Foundation of China (No. 31830084, 31672336), and also supported by the construction funds for “Double First-Class” initiative for Nankai University (No.s of 96172158, 96173250 and 91822294).

## Supporting Information

### Du *et al*

#### SI text

##### Fossil Record of Coreidae

All the confirmed fossil records of Coreidae during Mesozoic are preserved in China, the oldest of which is from the Upper Triassic strata (1–3). Fossil representatives are relatively common in Tertiary strata, being known from Eocene strata of the United States (4, 5); Oligocene rocks of France (6), Germany (7, 8); Miocene strata of China (9–13), Croatia (14) and Germany (14); and Pliocene rocks of France (15).

##### Taxonomic status of the nymph specimen preserved in amber

The extremely expanded and oversized antennae distinguish the new coreid from all other previously known fossil and extant species. In addition to Coreinae, Coreidae includes three other subfamilies. We can rule the nymph out of them by the following characteristics: Hydarinae are recognized by possession of a third antennal segment that is more than twice as long as the second segment, anterior and posterior lobes of metathoracic peritreme that are completely separated (16); Meropachyinae are characterized by a distal tooth or spine on the hind tibia, a curved and usually strongly incrassate hind femur (17); Pseudophloeinae are distinguished by a granulated surface of the pronotum, scutellum and hemelytra, with each granule bearing small adpressed setae (18). All previous fossil examples of Coreidae have been reported from rock impressions, and are generally poorly preserved. *Yuripopovina magnifica* of family Yuripopovinidae is the oldest known specimen of the superfamily Coreoidea preserved in the Cretaceous Lebanese amber (19).

The coreid nymph specimen preserved in amber described in the present study most likely belongs to the subfamily Coreinae owing to the following features: third antennal segment only slightly longer than second segment; hind tibia not curved and without prominent tooth or spine distally; and pronotum smooth and not granular. This is the first report of a leaf-footed bug in amber and the second representative of the Coreoidea preserved in amber.

##### Systematic paleontology

Order Hemiptera Linnaeus, 1758

Suborder Heteroptera Latreille, 1810

Infraorder Pentatomomorpha Leston, Pendergrast and Southwood, 1955

Superfamily Coreoidea Reuter, 1815

Family Coreidae Leach, 1815

Subfamily Coreinae Leach, 1815

Genus *Magnusantenna* Du & Chen gen. nov.

##### *Magnusantenna wuae* Du & Chen gen. et sp. nov

(Figs. 1, S1–3)

##### Type species

*Magnusantenna wuae* Du & Chen gen. et sp. nov.

##### Remarks

*Magnusantenna* Du & Chen gen. nov. is similar to the extant *Chariesterus* Laporte, 1832 in the following ways: body slender; lateral margins parallel; head subquadrate; compound eyes prominent and protruding; antennal socket protruding forward; antennae subequal to the length of the body; third antennal segment variously foliate; pronotum narrowed anteriorly, without collar; and hind tibiae not expanded. However, *Magnusantenna* gen. nov. can be distinguished from *Chariesterus* by several characteristics: first antennal segment slightly fusiform; second antennal segment approximately rectangular and spreading; fourth antennal segment exhibiting very large triangular spread; rostrum segments each of different length; and pronotum without spinose humeri. Conversely, *Chariesterus* exhibits several characteristics that differ from those of *Magnusantenna* gen. nov.: first antennal segment somewhat triquetral, usually bearing small denticles or acute spines, slightly curved at least in the basal area; second and fourth antennal segments not expanded; and rostrum segments subequal in length, diverging posteriorly to form prominent spinose humeri (20, 21). *Magnusantenna* gen. nov. is markedly different from all previously described fossil Coreidae in the scale of the antennal exaggeration.

##### *Magnusantenna wuae* Du & Chen gen. et sp. nov

(Figs. 1, S1–3)

##### Diagnosis

as for genus.

##### Description

Body slender, length 6.67 mm, width 0.76 mm (Figs. S1, S3). Head subquadrate, length 0.55 mm, width 0.56 mm (Figs. 1, S1, S3).

Labrum long triangle, basal area slightly broad, gradually narrowing toward apex (Figs. 1B, S1, S3). Compound eyes large and spherical, located on the center of the lateral margins of the head and protruding outward significantly (Figs. 1B, S1, S3). Rostrum with four segments; first segment close to the ventral surface of the head, reaching the anterior margin of the compound eye, length 0.47 mm; second segment vertical to the body, longest rostrum segment, length 0.92 mm; third and fourth segments parallel to the body, pointing forward, length 0.79 mm and 0.42 mm, respectively, and the apex of the fourth segment sharp (Fig. 1B).

Antennae nearly 12.3 times longer than the head and 4.4 times wider than the head. Antennal socket robust, extending in front of the head. Antennae with four segments, length 6.78 mm, slight longer than body length, with significant expansion except for the first segment (Figs. 1A, S1, S3). First antennal segment inflated, 0.42 mm long and 0.20 mm wide (Figs. 1C, S1, S3). Second antennal segment approximately rectangular and expanding, with a few setae on the surface, lateral margins serrated and setaceous, distal margin of antennal axis with a sharp angle at apex, proximal margin of antennal axis thickened and cutinized, segment length 1.88 mm and width 0.29 mm (Figs. 1D, S1, S3). Third antennal segment petal-shaped, 2.17 mm long and 1.14 mm wide; basal area obtusely rounded, middle of apical area with a sharp angle, all margins bear minute setae; distal expansion of antennal axis with sparse setae on the surface, proximal expansion of antennal axis with dense setae on the surface, proximal margin with strong keratin thickening (Figs. 1E, S1, S3). Fourth antennal segment triangular, 3.15 mm long and 2.49 mm wide, basal area obtusely rounded, apical area a long arc, all margins with minute setae (Figs. 1F, S1, S3); distal expansion of antennal axis with sparse setae on the surface (Fig. 1G), proximal expansion of the antennal axis with dense setae on the surface (Fig. 1H), proximal margin with strong keratin thickening (Fig. 1I).

Pronotum trapeziform, center with shallow longitudinal groove, length 0.65 mm and width 0.57 mm (Figs. S1–3). Mesonotum trapeziform, center with shallow longitudinal groove, length 0.49 mm and width 0.58 mm; lateral margin bearing forewing bud, long ovoid, length 0.65 mm and width 0.21 mm, basal area narrow, apical area narrowly rounded, posterior margin surpassing anterior margin of the metanotum, overlapping with the basal area of the hindwing bud (Figs. S1–3). Metanotum transversely wide, anterior margin nearly fused with posterior margin of mesonotum, length 0.39 mm and width 0.67 mm, lateral margin with hindwing bud, basal area wide, apical area narrowly rounded, posterior margin not reaching the anterior margin of the first abdominal tergite, length 0.34 mm and width 0.24 mm (Figs. S1–3).

Fore femora cylindrical and slightly thick, length 1.98 mm. Fore tibiae narrower than the femora, length 2.03 mm. Fore tarsi with two segments, apices with two claws, length 0.71 mm (Figs. S1–3). Middle femora slightly shorter than the fore femora, cylindrical, length 1.52 mm. Middle tibiae narrower than the femora, length 1.75 mm. Middle tarsi with two segments, apices with two claws, length 0.67 mm (Figs. S1–3). Hind femora long and thick, cylindrical, length 1.69 mm. Hind tibiae narrower than the femora, length 2.46 mm. Hind tarsi with two segments, apices with two claws, length 0.68 mm.

Abdomen length 4.35 mm, width 0.61 mm, nine visible segments. First and second abdominal tergite transversally wide. From third to eighth segment, abdominal tergites longer than the first two tergites. Black scent gland markings visible between the third and fourth, fourth and fifth, and fifth and sixth abdominal tergites. Ninth abdominal tergite trapezoidal, basal area wide, apical area slightly narrow, no recognizable genital structure that should be present in the abdomen (Figs. S1–3).

##### Remarks

*M. wuae* Du & Chen gen. et sp. nov. resembles the extant coreid *Chariesterus antennator* (Fabricius, 1803) (23). In addition to the similarities and differences documented in the Remarks section for the genus, the third antennal segments of both species are obovately dilated, with a width more than one-third of the length of the segment and the apex with an obvious angle. However, *M. wuae* gen. et sp. nov. does not have a notch at the apex of the third antennal segment, and the length ratio of each antennal segment from the first to the fourth is 42:188:217:315. In contrast, *C. antennator* has a conspicuous notch at the apex of the third antennal segment, and the length ratio of each antennal segment from the first to the fourth is 105:83:60:51 (21, 24). Therefore, *M. wuae* gen. et sp. nov. is sufficiently distinct from *C. antennator* to justify erection of a new genus and species.

#### SI Figures

**Fig. S1.**
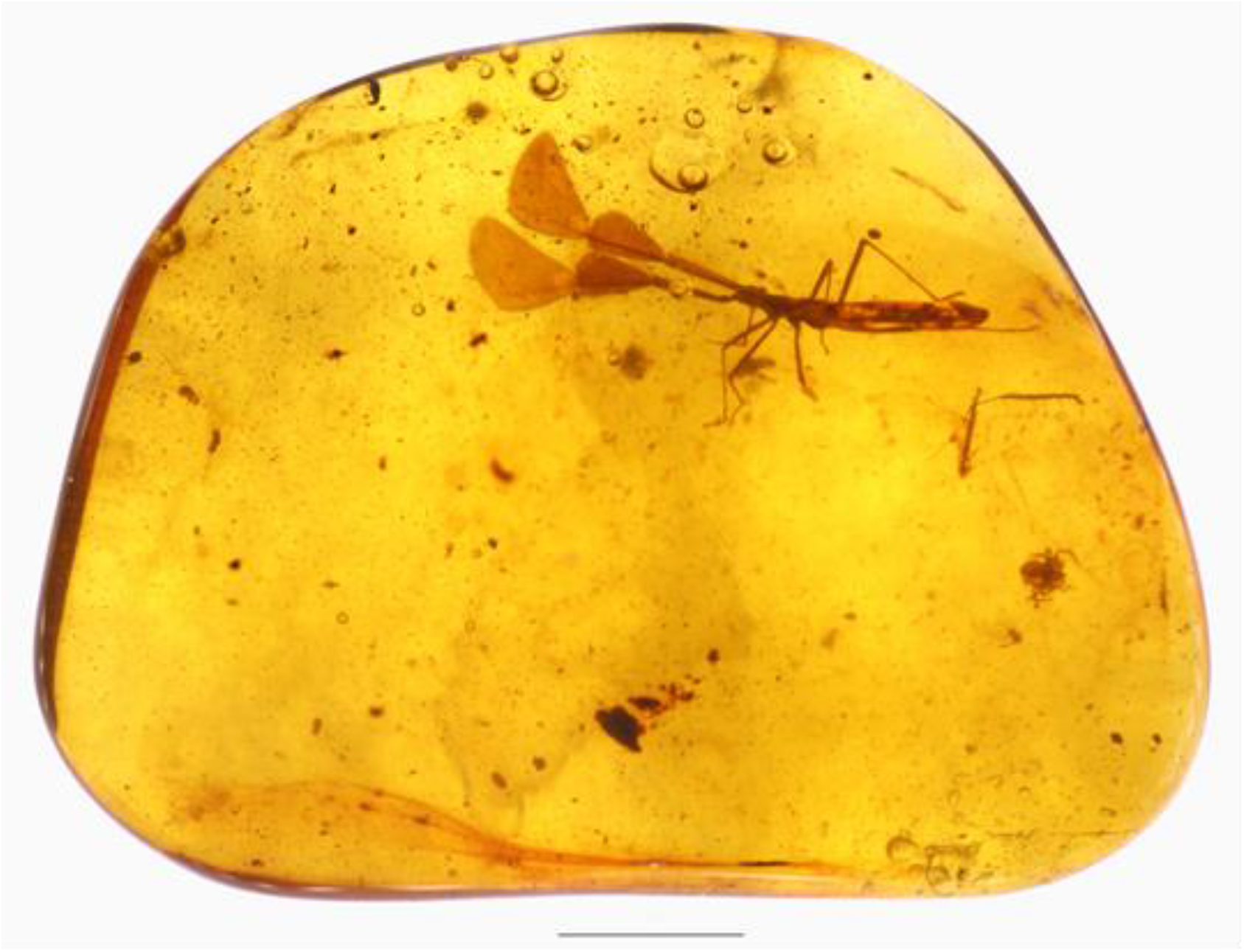
Overview of the amber. Scale bar, 5 mm.

**Fig. S2.**
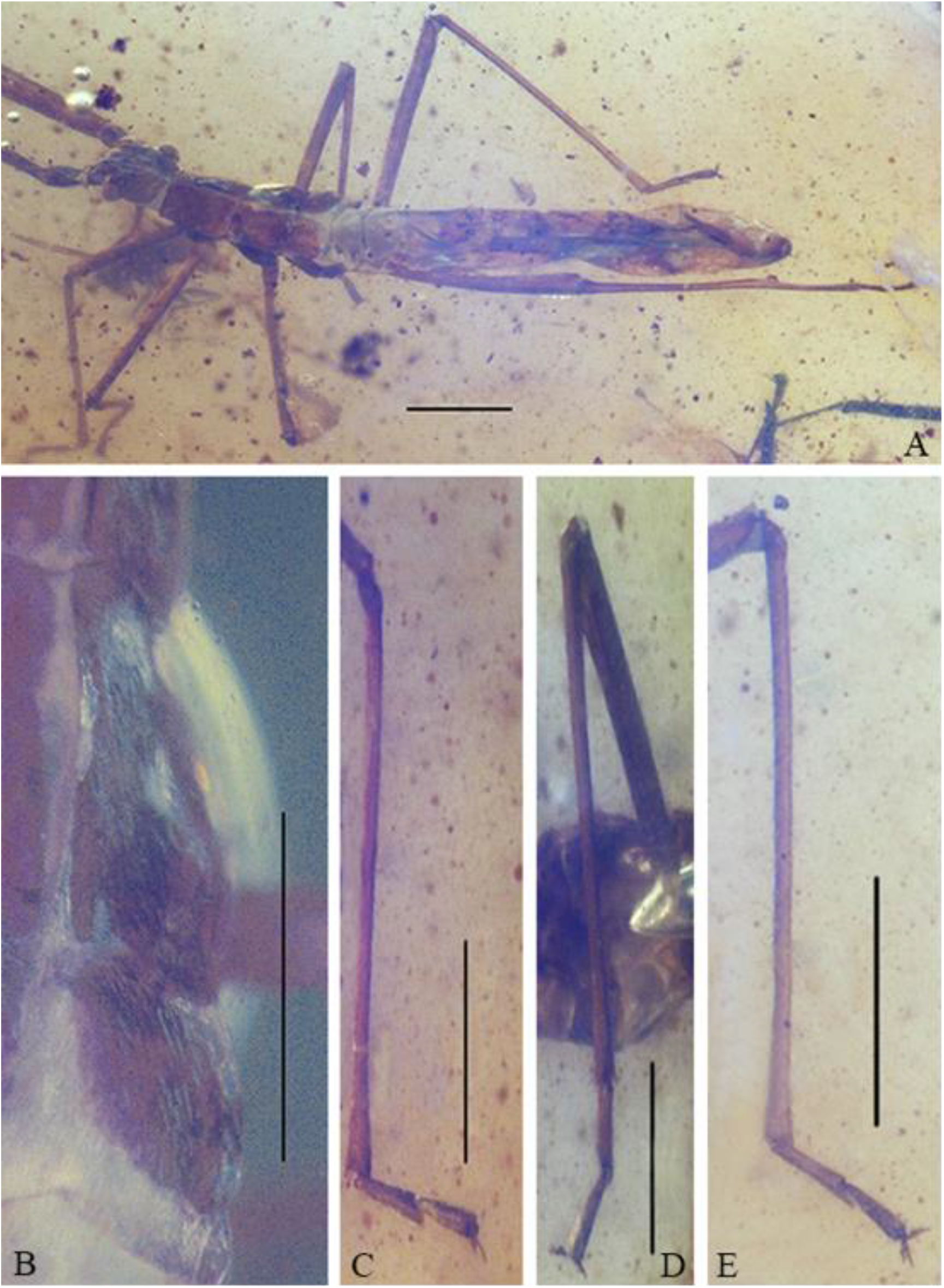
Thorax and abdomen of *M. wuae* gen. et sp. nov. (STJS0003). (A) Dorsal view of body. Scale bar, 1 mm. (B) Dorsal view of wing buds. Scale bar, 500 μm. (C) Lateral views of tibiae and tarsi of fore leg. Scale bar, 1 mm. (D) Lateral views of tibiae and tarsi of middle leg. Scale bar, 500 μm. (E) Lateral views of tibiae and tarsi of hind leg. Scale bar, 1 mm.

**Fig. S3.**
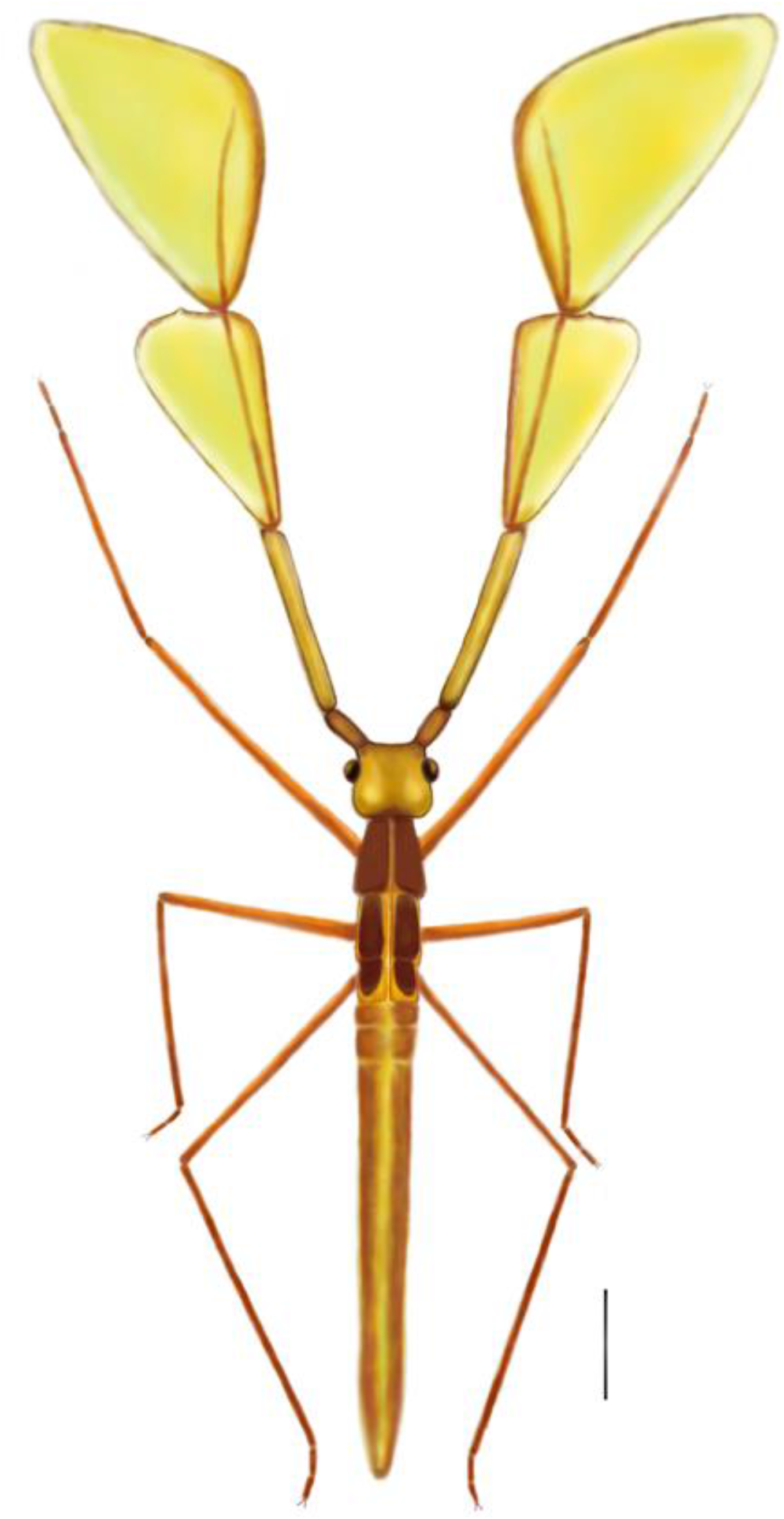
Reconstruction of the habitus of *M. wuae* gen. et sp. nov. (STJS0003). Scale bar, 1 mm.

**Fig. S4.**
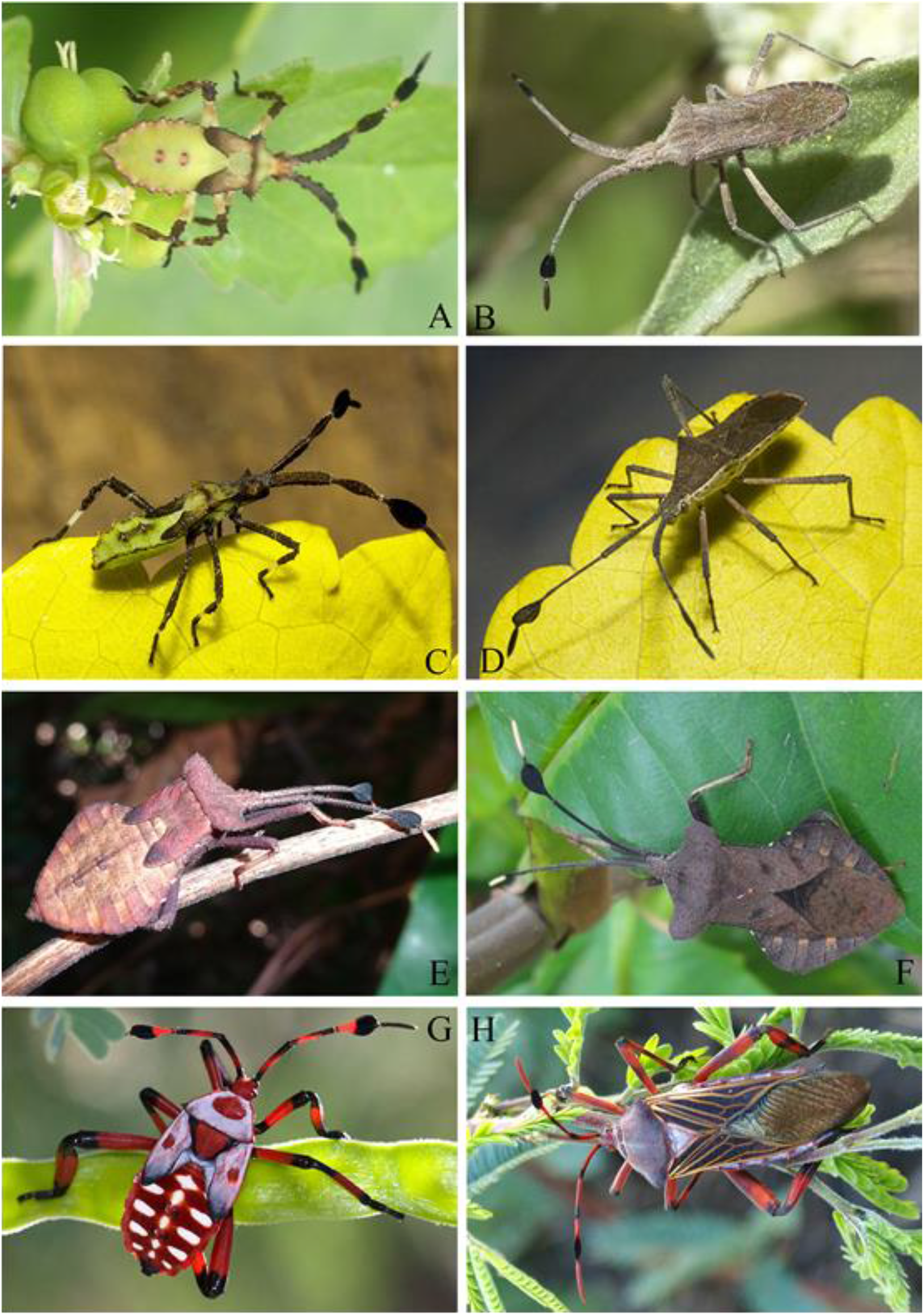
Images of some modern coreids. (A–B) *Chariesterus antennator* (A. Courtesy of Ms. Melanie Schuchart, downloaded from https://www.inaturalist.org/photos/20221697; B. Courtesy of Mr. Steve Kerr, downloaded from http://www.inaturalist.org/photos/1758620). (C–D) *Chariesterus armatus* (Courtesy of Mr. Branco, downloaded from https://www.flickr.com/photos/brutamonte). (E–F) *Dalader* sp. (Courtesy of Kwan downloaded from http://www.natureloveyou.sg). (G–H) *Thasus neocalifornicus* (Dr. Margarethe Brummermann, downloaded from http://arizonabeetlesbugsbirdsandmore.blogspot.com). A, C, E, G. Nymphs; B, D, F, H. Adults.

## References

1. J. A. C. Uy, G. Borgia, Sexual selection drives rapid divergence in bowerbird display traits. Evolution 54, 273–278 (2000).

2. N. A. Melnycky, R. B. Weladji, Ø. Holand, M. Nieminen, Scaling of antler size in reindeer (*Rangifer tarandus):* sexual dimorphism and variability in resource allocation. J. Mammal. 94, 1371–1379 (2013).

3. D. J. Emlen, The evolution of animal weapons. Annual Review of Ecology Evolution & Systematics 39, 387–413 (2008).

4. D. J. Emlen, L. Corley Lavine, B. Ewen-Campen, On the origin and evolutionary diversification of beetle horns. Proc. Natl. Acad. Sci. U.S.A. 104, 8661–8668 (2007).

5. E. L. McCullough, B. W. Tobalske, D. J. Emlen, Structural adaptations to diverse fighting styles in sexually selected weapons. Proc. Natl. Acad. Sci. U.S.A 111, 14484–14488 (2014).

6. T. L. Johnson, M. R. E. Symonds, M. A. Elgar, Sexual selection on receptor organ traits: younger females attract males with longer antennae. Sci. Nat. 104, 44 (2017).

7. C. Darwin, The descent of man, and selection in relation to sex (William Clowes and Sons, 1871), vol. 1, pp. 253.

8. G. I. Holwell, K. L. Barry, M. E. Herberstein, Mate location, antennal morphology, and ecology in two praying mantids (Insecta: Mantodea). Biol. J. Linn. Soc. 91, 307–313 (2007).

9. M. A. Elgar, T. L. Johnson, M. R. E. Symonds, Sexual selection and organs of sense: Darwin’s neglected insight. Anim. Biol. 69, 63–82 (2019).

10. M. Nakano, M. Morgan-Richards, A. J. R. Godfrey, A. C. McCormick, Parthenogenetic females of the stick insect *Clitarchus hookeri* maintain sexual traits. Insects 10, 1–16 (2019).

11. Q. K. Wang et al., Antennal scales improve signal detection efficiency in moths. Proceedings of the Royal Society B-Biological Sciences 285, 20172832 (2018).

12. D. T. Gwynne, W. J. Bailey, Female-female competition in katydids: sexual selection for increased sensitivity to male signals? Evolution 53, 546–551 (1999).

13. X. Z. Yan, C. P. Deng, X. J. Sun, C. Hao, Effects of various degrees of antennal ablation on mating and oviposition preferences of the diamondback moth, *Plutella xylostella* L. Journal of Integrative Agriculture 13, 1311–1319 (2014).

14. A. Jayaweera, K. L. Barry, Male antenna morphology and its effect on scramble competition in false garden mantids. The Science of Nature 104, 75 (2017).

15. L. Krogmann, M. S. Engel, G. Bechly, A. Nel, Lower Cretaceous origin of long-distance mate finding behaviour in Hymenoptera (Insecta). J. Syst. Palaeontol. 11, 83–89 (2013).

16. T. Bao, B. Wang, J. Li, D. Dilcher, Pollination of Cretaceous flowers. Proc. Natl. Acad. Sci. U.S.A. 116, 24707–24711 (2019).

17. S. Wedmann, S. Bradler, J. Rust, The first fossil leaf insect: 47 million years of specialized cryptic morphology and behavior. Proc. Natl. Acad. Sci. U.S.A. 104, 565–569 (2007).

18. C. Y. Cai et al., Early origin of parental care in Mesozoic carrion beetles. Proc. Natl. Acad. Sci. U.S.A. 111, 14170–14174 (2014).

19. P. D. L. Fuente, M. S. Engel, Early evolution and ecology of camouflage in insects. Proc. Natl. Acad. Sci. U.S.A. 109, 21414–21419 (2012).

20. T. Gao et al., Convergent evolution of ramified antennae in insect lineages from the Early Cretaceous of Northeastern China. Proc. R. Soc. B. 283, 20161448 (2016).

21. W. Wichard, A remarkable caddisfly with bipectinate antennae in Cretaceous Burmese amber (Insecta, Trichoptera). Cretaceous. Res. 69, 198–203 (2017).

22. Q. Liu et al., High niche diversity in Mesozoic pollinating lacewings. Nat. Commun. 9, 3793 (2018).

23. S. Du, Z. Hu, Y. Yao, D. Ren, New genus and species of the Yuripopovinidae (Pentatomomorpha: Coreoidea) from mid-Cretaceous Burmese amber. Cretaceous. Res. 94, 141–146 (2018).

24. T. Wappler, E. Guilbert, C. C. Labandeira, T. Hörnschemeyer, S. Wedmann, Morphological and behavioral convergence in extinct and extant bugs: the systematics and biology of a new unusual fossil lace bug from the Eocene. PLoS ONE 10, e0133330 (2015).

25. E. Barrera, H. Brailovsky, Descripcion de cuatro especies y una subespecie nuevas de la tribu anisoscelidini (Hemiptera-Heteroptera-Coreidae). Anales del Instituto de Biología Universidad Nacional Autónomia de México (Serie Zoología) 65, 45–62 (1994).

26. H. E. Hamouly, R. F. Sawaby, H. H. Fadl, Taxonomic review of the subfamily Pseudophloeinae (Hemiptera: Coreidae) from Egypt. Egypt J. Biol. 12, 108–124 (2010).

27. J. A. M. Fernandes, P. L. Mitchell, L. Livermore, M. Nikunlassi, “Leaf-footed bugs (Coreidae)” in True bugs (Heteroptera) of the neotropics, A. R. Panizzi, J. Grazia, Eds. (Springer Netherlands, 2015), pp. 549–605.

28. W. G. Eberhard, Sexual behavior of *Acanthocephala declivis guatemalana* (Hemiptera: Coreidae) and the allometric scaling of their modified hind legs. Ann. Entomol. Soc. Am. 91, 863–871 (1998).

29. R. T. Schuh, J. A. Slater, “Coreidae” in True bugs of the world (Hemiptera: Heteroptera): classification and natural history, R. T. Schuh, J. A. Slater, Eds. (Cornell University Press, 1995), vol. 89, pp. 274–279.

30. Paleobiology Database (2018) The Paleobiology Database. Checklist dataset https://doi.org/10.15468/zzoyxi accessed via GBIF.org on 2020-05-12.

31. G. Shi et al., Age constraint on Burmese amber based on U–Pb dating of zircons. Cretaceous. Res. 37, 155–163 (2012).

32. K. L. Prudic, K. Noge, J. X. Becerra, Adults and nymphs do not smell the same: the different defensive compounds of the giant mesquite bug (*Thasus neocalifornicus:* Coreidae). J. Chem. Ecol. 34, 734–741 (2008).

33. A. Ramsey et al., Towards an understanding of molecule capture by the antennae of male beetles belonging to the genus *Rhipicera* (Coleoptera, Rhipiceridae). The anatomical record 298, 1519–1534 (2015).

34. R. Mankin, M. Mayer, The insect antenna is not a molecular sieve. Experientia 40, 1251–1252 (1984).

35. S. Pekár, M. Hrušková, How granivorous *Coreus marginatus* (Heteroptera: Coreidae) recognises its food. Acta Ethol. 9, 26–30 (2006).

36. M. Elgar et al., Insect antennal morphology: the evolution of diverse solutions to odorant perception. The Yale journal of biology and medicine 91, 457–469 (2018).

37. D. K. McLain, L. B. Burnette, D. A. Deeds, Within season variation in the intensity of sexual selection on body size in the bug *Margus obscurator* (Hemiptera Coreidae). Ethol. Ecol. Evol. 5, 75–86 (1993).

38. Y. Wang et al., Ancient pinnate leaf mimesis among lacewings. Proc. Natl. Acad. Sci. U.S.A. 107, 16212–16215 (2010).

39. Y. Wang et al., Jurassic mimicry between a hangingfly and a ginkgo from China. Proc. Natl. Acad. Sci. U.S.A. 109, 20514–20519 (2012).

40. X. Y. Liu et al., Liverwort mimesis in a Cretaceous lacewing larva. Curr. Biol. 28, 1–7 (2018).

41. M. J. F. Martins, T. M. Puckett, R. Lockwood, J. P. Swaddle, G. Hunt, High male sexual investment as a driver of extinction in fossil ostracods. Nature 556, 366–369 (2018).

42. E. Morrow, C. Fricke, Sexual selection and the risk of extinction in mammals. Proc. R. Soc. Lond. B. 271, 2395–2401 (2004).

43. Y. Tanaka, Sexual selection enhances population extinction in a changing environment. J. Theor. Biol. 180, 197–206 (1996).

44. P. F. Doherty et al., Sexual selection affects local extinction and turnover in bird communities. Proc. Natl. Acad. Sci. U.S.A. 100, 5858–5862 (2003).

45. J. Bro-Jørgensen, Will their armaments be their downfall? Large horn size increases extinction risk in bovids. Anim. Conserv. 17, 80–87 (2014).

46. L. Xing, K. Niu, S. E. Evans, Inter-amphibian predation in the Early Cretaceous of China. Sci. Rep. 9, 7751 (2019).

47. S. R. Schachat, C. C. Labandeira, M. E. Clapham, J. L. Payne, A Cretaceous peak in family-level insect diversity estimated with mark–recapture methodology. Proc. R. Soc. B. 286, 20192054 (2019).

48. D. Zheng et al., Extreme adaptations for probable visual courtship behaviour in a Cretaceous dancing damselfly. Sci. Rep. 7, 44932 (2017).

49. Q. Wang, C. Shih, D. Ren, The earliest case of extreme sexual display with exaggerated male organs by two middle Jurassic Mecopterans. PLoS ONE 8, e71378 (2013).

50. R. D. Cruickshank, K. Ko, Geology of an amber locality in the Hukawng Valley, Northern Myanmar. J. Asian Earth Sci. 21, 441–455 (2003).

51. D. A. Grimaldi, M. S. Engel, P. C. Nascimbene, Fossiliferous Cretaceous amber from Myanmar (Burma): Its rediscovery, biotic diversity, and paleontological significance. Am. Mus. Novit. 62, 1–71 (2002).

## References

1. Q. B. Lin, Late Triassic insect fauna from Toksun, Xinjiang. Acta Palaeontol. Sin. 31, 313–335 (1992).

2. Y. C. Hong, Insecta. Palaeontological Atlas of North China, II, Mesozoic, 128–185 (1984).

3. Y. Hong, The study of Early Cretaceous insects of Kezuo, west Liaoning. Professional Papers of Stratigraphy & Palaeontology 18, 76–87 (1987).

4. S. H. Scudder, The Tertiary insects of North America (George Mason University Press, 1890), vol. 13, pp. 1–734.

5. T. D. A. Cockerell, Fossil insects from Colorado. The Entomologist 42, 170–174 (1909).

6. C. v. Heyden, Fossile Insekten aus der Braunkohle von Salzhausen. Palaeontographica 5, 115–120 (1858).

7. G. Statz, E. Wagner, Geocorisae (Landwanzen) aus den Oberoligocäner Ablagerungen von Rott. Journal of Differential Equations 34, 496–522 (1950).

8. Y. C. Hong, J. Cora, N. Johnson, Fossil insects in the diatoms of Shanwang. Bulletin of the Tianjin Institute of Geology and Mineral Resources 8, 1–15 (1983).

9. Y. C. Hong, W. L. Wang, Miocene Heteroptera and Coleoptera (Insecta) from Shanwang of Shandong Province, China. Journal of the Lanzhou University of Natural Science 33, 116–124 (1987).

10. J. F. Zhang, Miocene insects from Shanwang of Shandong, China and their bearing on palaeoenvironment. Proceedings of International Symposium on Pacific Neogene Continental and Marine Events, 149–156 (1989).

11. J. F. Zhang, X. Y. Zhang, Fossil insects of cicada (Homoptera) and true bugs (Heteroptera) from Shanwang, Shandong. Acta Palaeontol. Sin. 29, 337–348 (1990).

12. J. F. Zhang, B. Sun, X. Y. Zhang, Miocene insects and spiders from Shanwang, Shandong (Science Press, 1994), pp. 1–298.

13. O. Heer, Die Insektenfauna der Tertiärgebilde von Oeningen und von Radoboj in Croatien. Dritte Theil: Rhynchoten (Neue Denkschriften der allgemeinen Schweizerischen Gesellschaft für die gesammten Naturwissenschaften Zürich, 1853), pp. 1–138.

14. L. E. Piton, La faune entomologique des gisements mio-pliocenes du Massif Central. Revue des Sciences Naturelles d’Auvergne (N. S.) 1, 65–104 (1935).

15. H. Brailovsky, New genus and new species of Hydarini (Hemiptera, Heteroptera, Coreidae) from South America. Dtsch. Entomol. Z. 57, 85–88 (2010).

16. H. Brailovsky, E. Barrera, New species of *Merocoris* (*Merocoris*) Perty from Brazil, with keys to known subgenera and species of the tribe Merocorini (Hemiptera: Heteroptera: Coreidae: Meropachyinae). Fla. Entomol 92, 134–138 (2009).

17. H. E. Hamouly, R. F. Sawaby, H. H. Fadl, Taxonomic review of the subfamily Pseudophloeinae (Hemiptera: Coreidae) from Egypt. Egypt J. Biol. 12, 108–124 (2010).

18. D. Azar, A. Nel, M. Engel, R. Garrouste, A. Matocq, A new family of Coreoidea from the Lower Cretaceous Lebanese amber (Hemiptera: Pentatomomorpha). Polish Journal of Entomology 80, 627–644 (2011).

19. J. A. M. Fernandes, P. L. Mitchell, L. Livermore, M. Nikunlassi, “Leaf-footed bugs (Coreidae)” in True bugs (Heteroptera) of the neotropics, A. R. Panizzi, J. Grazia, Eds. (Springer Netherlands, 2015), pp. 549–605.

20. H. Ruckes, The genus *Chariesterus* de Laporte (Heteroptera, Coreidae). American Museum novitates 1721, 1–16 (1955).

21. J. C. Fabricius, “Coreus” in Systema rhyngotorum: secundum ordines, genera, species: adiectis synonymis, locis, observationibus, descriptionibus, J. C. Fabricius, Ed. (C. Reichard, 1803), pp. 191–202.

22. S. B. Fracker, *Chariesterus* and its neotropical relatives (Coreidae Heteroptera). Ann. Entomol. Soc. Am. 12, 227–230 (1919).

